# A Connectome Wide Functional Signature of Transdiagnostic Risk for Mental Illness

**DOI:** 10.1101/196220

**Authors:** Maxwell L. Elliott, Adrienne Romer, Annchen R. Knodt, Ahmad R. Hariri

## Abstract

**Background:** High rates of comorbidity, shared risk, and overlapping therapeutic mechanisms have led psychopathology research towards transdiagnostic dimensional investigations of clustered symptoms. One influential framework accounts for these transdiagnostic phenomena through a single general factor, sometimes referred to as the ‘p’ factor, associated with risk for all common forms of mental illness.

**Methods:** Here we build on past research identifying unique structural neural correlates of the p factor by conducting a data-driven analysis of connectome wide intrinsic functional connectivity (n = 605).

**Results:** We demonstrate that higher p factor scores and associated risk for common mental illness maps onto hyper-connectivity between visual association cortex and both frontoparietal and default mode networks.

**Conclusions:** These results provide initial evidence that the transdiagnostic risk for common forms of mental illness is associated with patterns of inefficient connectome wide intrinsic connectivity between visual association cortex and networks supporting executive control and self-referential processes, networks which are often impaired across categorical disorders.

## Introduction

Emerging research has identified a general factor of psychopathology that accounts for shared risk among internalizing, externalizing, and thought disorders across diverse samples(1–6). In contrast to the traditional clinical science model which compares cases of individuals meeting criteria for a categorical disorder to those not meeting these criteria (i.e., “healthy” controls), this general psychopathology or ‘p’ factor reflects an individuals’ latent liability for mental illness(7). For individuals with clinical disorders, higher p factor scores portend greater chronicity and symptom severity(1, 7). In “healthy” individuals, higher p factor scores reflect relative risk for developing future clinical disorder. Moreover, the p factor provides a framework for explaining the high rates of comorbidity as well as the shared genetic variance among categorical mental disorders(8, 9). As such, the p factor represents a potentially useful avenue for better understanding the shared and unique etiology of common mental illness. However, the biological mechanisms through which the p factor confers general risk for psychopathology remain unclear. Identifying such mechanisms is necessary for effectively leveraging the p factor to derive novel targets for clinical intervention and prevention.

Clinical neuroscience has begun to adopt transdiagnostic methodologies to accelerate the search for common neurobiological abnormalities across disorders(10). For example, a recent large meta-analysis of six categorical disorders reported a shared pattern of reduced gray matter volume in a distributed network supporting attention and cognitive control(11). In addition, we have recently examined the structural neural correlates of the p factor specifically(12). In our work, higher p factor scores and thus risk for common mental illness was associated with reduced gray matter volumes in the occipital lobes and neocerebellum. Furthermore, higher p factor scores were associated with reduced fractional anisotropy in pontine pathways linking the neocerebellum with the thalamus and prefrontal cortex. This network of brain regions is thought to be a forward monitor of incoming sensory information, generating and updating internal models for motor as well as cognitive tasks(13). In addition, activation of the neocerebellum has been associated with cognitive control tasks,(14) reflecting its contribution to the extended cognitive control network including the dorsolateral and medial prefrontal cortex(15). Thus, our observed p factor associations along with meta-analytic results suggest that transdiagnostic risk for common forms of mental illness may be associated with structural deficits in a network of brain regions supporting cognitive control. However, the putative functional consequences of these observed structural associations have not yet been examined.

Resting-state functional connectivity is a powerful tool in clinical neuroscience because it can be readily administered across patient populations(16, 17), demonstrates trait-like stability(18) as well as moderate heritability(19, 20), and represents a powerful probe of the intrinsic architecture of neural networks that play a primary role in shaping task-based network activity and associated behaviors(21). In addition, altered intrinsic functional connectivity within the default mode network (DMN), and frontoparietal network (FPN), both of which are linked to higher order cognition, have been broadly linked to psychopathology across categorical disorders(22–24). Thus, resting-state measures of intrinsic network connectivity represent one avenue for extending the structural associations of the p factor to variability in functional neural dynamics representing mechanisms through which risk may emerge.

Here, we investigate intrinsic functional connectivity correlates of the p factor in a volunteer sample of 614 university students from the Duke Neurogenetics Study. While our previous research in this sample identified discrete structural correlates of the p factor in the occipital lobes, neocerebellum, and pons, we opted for a whole-brain exploratory analysis of intrinsic connectivity to capture functional differences beyond these regions and impose minimal assumptions about the nature of p factor associations in the brain. While there are many exploratory methods for investigating resting state functional connectivity, we performed a Connectome-Wide Association Study (CWAS)(25) of the p factor using multidimensional matrix regression (MDMR)(26). In contrast to traditional seed-based approaches, MDMR allows a search across the whole brain for multivariate connectivity patterns that vary with p factor scores, while at the same time making few assumptions about the data or expected effects. Unlike clustering(27) or independent components analysis,(28) MDMR does not require *a priori* estimates of the dimensionality of the data or choosing networks or connections of interest. In addition, MDMR does not require arbitrary decisions about thresholding matrices (as in many graph analysis techniques(29)), while retaining the advantages of interpretability and visualization of traditional seed based approaches. For these reasons, we conducted a CWAS to identify associations between p factor scores and intrinsic functional connectivity.

## Methods

### Participants

Data for this study come from the Duke Neurogenetics study (DNS), which was designed to allow for examination of predictive links between genes, brain, behavior and risk for mental illness among 18 to 22-year-old university students. DNS participants were recruited primarily from the Duke University student body via flyers and online postings. After successful completion of the DNS protocol, participants received financial compensation as well as a free 23andMe account. While all 1333 DNS participants completed mental health assessments and structural neuroimaging, resting-state fMRI was only collected on a subset due to revisions of the MRI protocol to accommodate two new task-fMRI scans which led to removal the resting-state scans from the protocol. Specifically, resting-state data was collected on 614 consecutive participants; therefore, this subsample is broadly representative of the entire DNS sample and does not suffer from further selection bias. All participants provided informed consent in accordance with the Duke University Medical Center Institutional Review Board guidelines before participation. All participants were in good general health and free of the following conditions, known to artifactually influence MRI data collection: (1) medical diagnoses of cancer, stroke, head injury with loss of consciousness, untreated migraine headaches, diabetes requiring insulin treatment, chronic kidney or liver disease; (2) use of psychotropic, glucocorticoid or hypolipidemic medication; and (3) conditions affecting cerebral blood flow and metabolism (e.g., hypertension). One goal of the DNS was to study mental health and illness; therefore, participants were not excluded if they met criteria for substance abuse or a mental illness.

### Clinical Diagnosis

Current and lifetime DSM-IV Axis I disorder or select Axis II disorders was assessed with the electronic Mini International Neuropsychiatric Interview(30) and Structured Clinical Interview for the DSM-IV subtests(31) respectively. Importantly, diagnosis wasn’t an exclusion criterion, as the DNS seeks to establish broad variability in multiple behavioral phenotypes related to psychopathology. Allowing for a broad spectrum of symptoms is particularly critical for accurately deriving p factor scores. Nevertheless, no participants were taking any psychoactive medication during or at least 14 days prior to their participation. Of the 605 participants with data included in our analyses, 133 individuals had at least one DSM-IV diagnosis, including 76 with alcohol use disorders, 24 with non-alcohol substance use disorders, 33 with major depression disorder, 26 with bipolar disorder, 7 with panic disorder (no agoraphobia), 9 with panic disorder including agoraphobia, 4 with social anxiety disorder, 8 with generalized anxiety disorder, 10 with obsessive compulsive disorder, and 7 with eating disorders. While this is a university-based convenience sample that is not representative of the broader population in intelligence or parental education (due to selective admissions criteria of Duke University), the sample is broadly representative of the general population in terms rates of mental illness(32).

### Derivation of p factor scores

In previous work(12), our group replicated the p factor in the DNS using confirmatory factor analysis of self-report and diagnostic interview measures of internalizing, externalizing, and thought disorder symptoms. These p factor scores were extracted using the standard regression method from those analyses and standardized to a mean of 100 (SD = 15), with higher scores indicating a greater propensity to experience all forms of psychiatric symptoms. Further details on the derivation of the p-factor scores can be found in the supplement.

### Image acquisition

Each participant was scanned using one of two identical research-dedicated GE MR750 3 T scanners equipped with high-power high-duty-cycle 50-mT/m gradients at 200 T/m/s slew rate, and an eight-channel head coil for parallel imaging at high bandwidth up to 1MHz at the Duke-UNC Brain Imaging and Analysis Center. A semi-automated high-order shimming program was used to ensure global field homogeneity. A series of 34 interleaved axial functional slices aligned with the anterior commissure-posterior commissure plane were acquired for full-brain coverage using an inverse-spiral pulse sequence to reduce susceptibility artifacts (TR/TE/flip angle=2000 ms/30 ms/60; FOV=240mm; 3.75×3.75×4mm voxels; interslice skip=0). Four initial radiofrequency excitations were performed (and discarded) to achieve steady-state equilibrium. For each participant, 2 back-to-back 4-minute 16-second resting state functional MRI scans were acquired. Participants were instructed to remain awake, with their eyes open during each resting state scan. To allow for spatial registration of each participant’s data T1-weighted images were obtained using a 3D Ax FSPGR BRAVO with the following parameters: TR = 8.148 ms; TE = 3.22 ms; 162 axial slices; flip angle, 12°; FOV, 240 mm; matrix =256×256; slice thickness = 1 mm with no gap; and total scan time = 4 min and 13 s.

### Image Processing

Anatomical images for each subject were skull-stripped, intensity-normalized, and nonlinearly warped to a study-specific average template in the standard stereotactic space of the Montreal Neurological Institute template using the ANTs SyN registration algorithm(33, 34). Time series images for each subject were despiked, slice-time-corrected, realigned to the first volume in the time series to correct for head motion using AFNI tools(35), coregistered to the anatomical image using FSL’s Boundary Based Registration(36), spatially normalized into MNI space using the non-linear ANTs SyN warp from the anatomical image, resampled to 2mm isotropic voxels, and smoothed to minimize noise and residual difference in gyral anatomy with a Gaussian filter set at 6-mm full-width at half-maximum. All transformations were concatenated so that a single interpolation was performed.

Time-series images for each participant were furthered processed to limit the influence of motion and other artifacts. Voxel-wise signal intensities were scaled to yield a time series mean of 100 for each voxel. Motion regressors were created using each subject’s 6 motion correction parameters (3 rotation and 3 translation) and their first derivatives(37, 38) yielding 12 motion regressors. White matter (WM) and cerebrospinal fluid (CSF) nuisance regressors were created using CompCorr(39). Images were bandpass filtered to retain frequencies between .008 and .1 Hz, and volumes exceeding 0.25mm frame-wise displacement or 1.55 standardized DVARS(40, 41) were censored. Nuisance regression, bandpass filtering and censoring for each time series was performed in a single processing step using AFNI’s 3dTproject. Participants were excluded if they had less than 185 TRs left after censoring (resulting in inadequate degrees of freedom to perform nuisance regressions), resulting in a final sample of 605 subjects.

### CWAS

To make the analysis computationally tractable, time-series were extracted from a parcellated atlas instead of using voxelwise data. We used the Lausanne atlas parcellated into 1015 equally sized regions through the program easy_lausanne (github.com/mattcieslak/easy_lausanne). Time-series data for each subject were then processed using CWAS. Described extensively elsewhere(25), CWAS consists of 3 processing steps. First, beginning with a single ROI time-series, seed-based connectivity analysis is conducted to generate a whole-brain functional connectivity map for each participant. Second, the average distance (1 minus the Pearson correlation) between each pair of participant’s functional connectivity maps is computed, resulting in a distance matrix encoding the multivariate similarity between each participant’s connectivity map. Finally, multi-dimensional matrix regression (MDMR) is used to generate a pseudo-F statistic quantifying the strength of the association between the phenotype of interest, here p factor scores, and the distance matrix created in the second step. The advantage of MDMR is allowing covariates to be entered into the regression and utilizing non-parametric permutation to generate p-values for each ROI. These three steps are repeated for each of the 1015 ROIs, resulting in a whole-brain map that represents the association between p factor scores and whole-brain connectivity at each ROI. CWAS was performed to identify seed regions with whole-brain patterns of connectivity are related to p factor scores. Participant sex was included as a covariate, and 500,000 permutations were performed to generate p-values. To minimize false positives across the 1,015 ROIs, a false discovery rate(42) (FDR) correction was applied. The threshold for statistical significance was set at q = .05.

### Seed-based analyses

MDMR identifies a set of ROIs with patterns of whole-brain connectivity associated with p factor scores. However, it is still unclear how the connectivity of these ROIs relates to the scores. Previous research using CWAS(25, 43, 44) has demonstrated the utility of using traditional seed-based connectivity follow-up analyses to better understand the networks and brain regions that drive the associations discovered through MDMR. Similar analyses were performed here for each ROI identified via MDMR. Seed-based connectivity maps were created and correlations were converted to Z statistics via the Fischer R to Z transform. Whole-brain correlations between these connectivity values and p factor scores were calculated, including sex as a covariate. Importantly, these follow-up analyses do not represent independent statistical tests as they were performed post-hoc to the family wise error controlled MDMR findings. Accordingly, these follow up analyses maps are not thresholded to visualize all information that was relevant to the MDMR step.

## Results

### Demographics

From the 614 participants who completed two resting-state scans, 605 had data that survived quality control procedures. Of these, 336 were women, and the mean age was 20.23±1.19 years old. Scores for the p factor ranged from 76.71to 191.96 with a mean of 99.80, sd of 15.39.

### Multi-dimensional matrix regression

Whole brain maps from 1,015 ROIs were compared to estimate the multivariate distance (dissimilarity) between each subject map at every ROI. MDMR was then used to statistically test the association between these distances and individual p factor scores. MDMR revealed that four ROIs had whole-brain connectivity patterns that were significantly associated with p factor scores. This included the left lingual gyrus(x = 28, y = 85, z = −18; corrected p = .9680), right middle occipital gyrus (x = −31, y = 94, z = −0; corrected p = .9743), and two adjacent parcels of the left middle occipital gyrus (x = 32, y = 93, z = −5; corrected p = .9949) and (x = 30, y = 96, z = 0; corrected p = .9949) (Figure 1).

**Figure 1.**
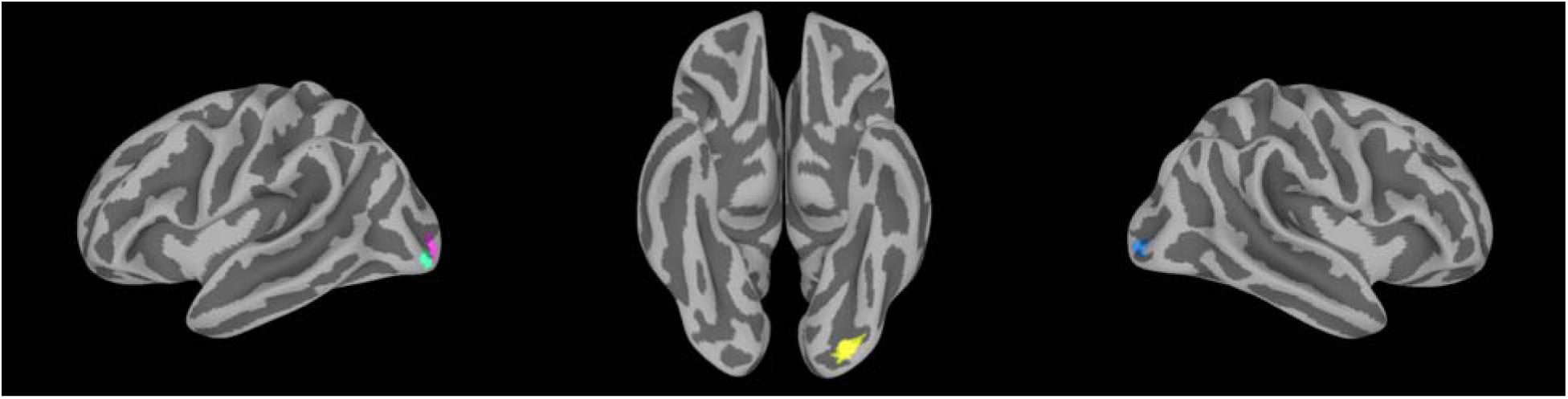
Data driven multi-dimensional matrix regression (MDMR) analysis revealed four regions with whole-brain connectivity patterns significantly associated with p factor scores: two adjacent parcels of the left middle occipital gyrus (left panel), left lingual gyrus (middle panel), and right middle occipital gyrus (right panel). These four clusters are projected onto a surface volume for visualization.

### Follow-up intrinsic connectivity analyses

The follow-up connectivity analyses of each seed identified through MDMR revealed the primary network associations for each seed as well as their pattern of whole-brain connectivity associated with p factor scores. These analyses showed striking convergence across MDMR-selected ROIs wherein the mean whole-brain pattern of connectivity for each seed showed subtle variation, but largely outlined the canonical resting-state visual processing network(45). The connectivity of each ROI with visual and somatosensory regions decreased with increasing p factor scores, while the connectivity between each ROI and transmodal association regions(46) increased with increasing p scores (Figure 2).

**Figure 2.**
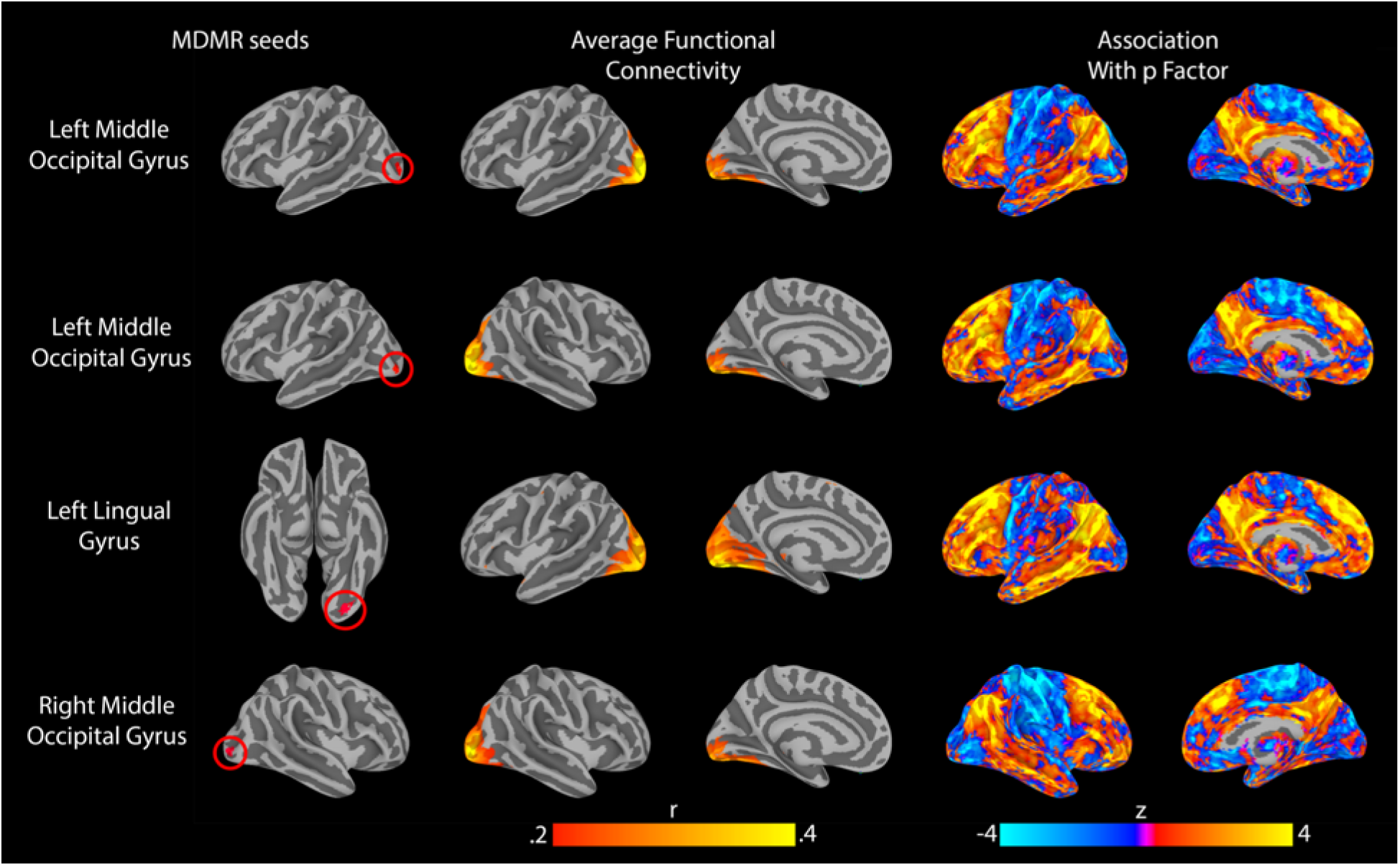
Follow-up connectivity analyses of the four seeds identified through MDMR revealed a highly-conserved pattern of altered connectivity between visual association cortex and both frontoparietal and default mode networks as a function of p factor scores. All results were projected from the volume onto a surface to aid visualization. Left panel: MDMR-derived seed regions. Middle panel: average intrinsic connectivity for each seed. Right panel: connectome wide intrinsic connectivity patterns for each seed as a function of p factor scores.

Further analyses were conducted to better characterize the above consistent patterns of p factor associations with the intrinsic connectivity of all seeds by averaging the independent whole-brain connectivity maps. The resulting average z-scores were summarized for each of the 7 Yeo networks(47) to quantify their respective contribution to the associations with p factor scores (Figure 3). These analyses revealed the DMN and FPN as the major networks for which intrinsic functional connectivity was positively correlated with p factor scores. In contrast, a more modest but notable negative correlation was observed between p factor scores and the intrinsic functional connectivity between the visual association cortex and somatomotor network.

**Figure 3.**
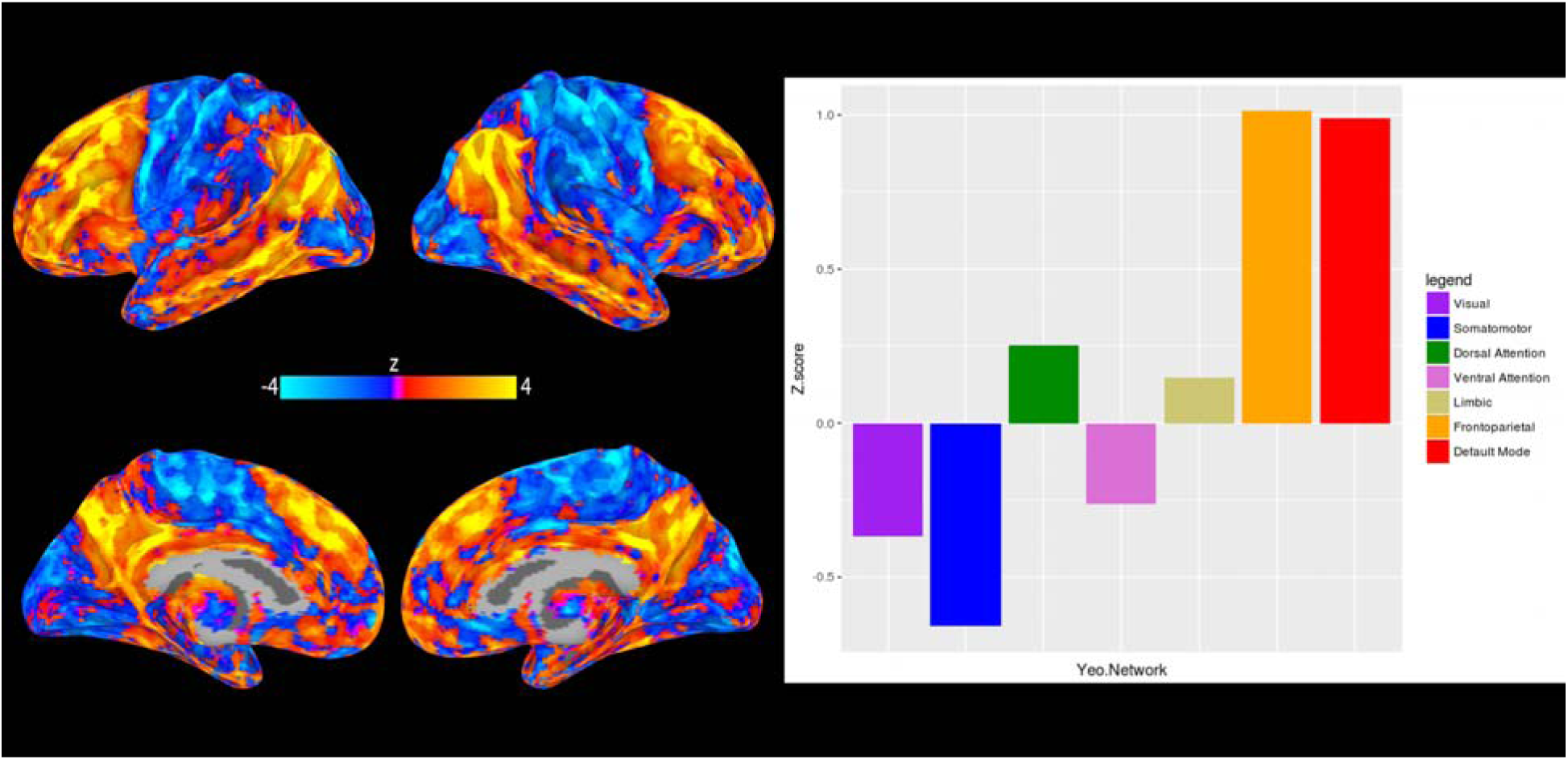
Mean pattern of intrinsic connectivity as a function of p factor scores across the networks associated with each of the four MDMR-derived seeds in visual association cortex (left panel). The relative contributions of seven canonical intrinsic cerebral networks(47) to this mean pattern of connectivity (right panel).

## Discussion

Here, we provide a novel extension of prior structural neural correlates of the p factor to the intrinsic architecture of the whole-brain functional connectome. Our unconstrained connectome-wide MDMR analysis revealed a circumscribed relationship between p factor scores and the whole-brain intrinsic connectivity of 4 nodes in visual association cortex. These findings are generally consistent with our earlier work finding a negative correlation between p factor scores and gray matter volume in the occipital cortex(12). Further investigation of the patterns of intrinsic connectivity driving this relationship primarily implicated hyper-connectivity between visual association cortex and heteromodal frontoparietal (FPN) and default mode networks (DMN). While differences in the intrinsic functional connectivity of visual areas is not commonly thought of as a core feature of psychopathology, our findings are not unique in pointing to dysfunction in visual association cortex and are consistent with a growing body of literature implicating sensory processing in transdiagnostic research.

Selection and suppression of incoming sensory information is an important component of goal directed behavior. Functional connectivity between visual and heteromodal association cortices (including FPN and DMN) has been shown to be critical for selecting task-relevant information(48, 49). Here we find that individual differences in the functional connectivity of visual association cortex with the FPN and DMN are associated with the p factor. Although speculative, our findings may indicate more effortful or less efficient integration of bottom-up sensory information with attentional demands and executive control processes in those at higher risk for mental illness. The specificity of this pattern to visual and not other sensory association cortices may reflect the dominance of the visual modality in guiding human perception of the external world and, possibly equally, the construction of internal models necessary for higher cognitive processes including executive control(50, 51).

Supporting evidence can be found in studies of schizophrenia and bipolar disorder, where visual network connectivity has been implicated in deficits involving the binding of visual objects(52) and in processing of visual stimuli(53). Functional connectivity between frontal association and visual cortex has also been associated with disrupted working memory in depression(54, 55) and in neurocognitive deficits in schizophrenia(56). While the visual cortices are not often thought of as primary to dysfunction in psychopathology, these studies suggest that visual cortical dysfunction may play a role in neurocognitive deficits present in many forms of psychopathology(57–59). Additionally, when assumptions are relaxed and whole-brain, resting-state connectivity analyses are performed, connections between the visual cortex and frontal association cortex have been shown to be predictive of psychopathology in depression(60) and schizophrenia(61). It is possible that the relative sparsity of links between visual cortex dysfunction and psychopathology partially reflects a bias in resting-state analyses towards strong assumptions about where in the brain findings are expected, which could result in missing associations with visual networks. Now that many large imaging datasets that include psychiatric data have been publicly released, our findings encourage further unbiased, data-driven whole-brain analyses in search of transdiagnostic neural correlates of psychopathology.

While our findings implicate visual association cortex in the general liability for mental illness, they do so primarily through its connectivity with the FPN and DMN. These networks consist of heteromodal association cortex that processes information from multiple sensory domains, and consist of brain regions most implicated in higher order thought and executive control of other networks(46). The unique role of the FPN and DMN in complex cognition(62–66) place them centrally in many etiologic theories of psychopathology(22, 23, 67–69), making their primary role in driving the association between visual cortex connectivity and p factor scores particularly relevant.

The frontoparietal network in particular has been linked to the core cognitive faculty of executive control,(21, 63, 68) which contributes to mental health and general well-being by shaping successful goal directed behavior(70). Fittingly, disrupted FPN activity has been linked to psychopathology across categorical disorders including schizophrenia(71), depression,(72) and bipolar disorder(73). Building off of this body of research, an emerging theory suggests that the relative integrity of the FPN and associated executive control mechanisms are fundamental for the capacity to self-regulate, manage symptoms, and succeed in treatment(22). Our current findings are consistent with this framework by demonstrating that higher p factor scores regardless of diagnosis are associated with relative hyper-connectivity of the FPN with the visual association cortex, suggesting one way through which FPN dysfunction may be manifest as psychopathology.

In addition to the frontoparietal network, our analyses implicate hyper-connectivity between the visual association cortex and default mode network as a function of higher p factor scores. The DMN has been generally linked to introspection, autobiographical memory, and future-oriented thought(69). Interestingly, DMN activity is suppressed in attention demanding tasks(69, 74) and altered DMN activity has been broadly observed across categorical psychiatric disorders(23, 67). Connectivity between the DMN and visual association cortex is important in the suppression of internally generated distracting information(49). Taken together, transdiagnostic risk for mental illness as indexed by p factor scores may lead to more effortful or less efficient processing when internally generated thought and externally generated sensory information compete for attention.

While providing initial evidence that broad risk for all forms of common mental illness is manifest as alterations in the intrinsic connectivity of functional neural networks, our analyses were exploratory by design and replication in independent samples is needed. Given prior research implicating the FPN and DMN across categorical disorders, we focused our above discussion on the potential relevance of intrinsic connectivity between visual association cortex and these networks in the emergence of transdiagnositic risk for mental illness. While the intrinsic connectivity of these networks also exhibited an outsized influence on the association with p factor scores, variation between visual association cortex and other resting-state networks contributed as well, albeit more modestly (Figure 3). MDMR uses information from all whole-brain connections in selecting seeds, and the inferential significance comes from the aggregate of connections rather than any one in particular. Thus, formally testing the relative contributions of different networks is not typically conducted. While we think future studies of the p factor will benefit from using our observations of intrinsic connectivity between visual association cortex and both DMN and FPN as *a priori* starting points, the potential relevance of other networks should not be ignored until the patterns reported herein are replicated.

Additional limitations, which can be addressed in future research, include the relatively limited range of psychopathology, especially severe forms including psychosis, represented in our volunteer sample of young adults. Future research should extend our analyses to more diverse populations including individuals with severe mental illness. While the DNS is broadly representative of population base rates of common forms of mental illness(32), it is not representative of the general population in terms of socioeconomics or intelligence. Thus, extension of these findings to population representative samples is needed. In addition, our results need to be replicated in well-powered independent samples to establish the reliability of these associations and provide unbiased estimates of the true effect sizes(75). In fact, we adopted a rigorous data-driven, unbiased approach in the current discovery analyses to minimize false positives and effect size inflation (i.e., “Winners Curse”) and bolster future attempts at replication. Our current analyses were also limited to the intrinsic connectivity of nodes within the cerebrum as our resting-state fMRI acquisition protocol did not afford full coverage of the cerebellum, including the neocerebellar subregion identified in our earlier structural analyses. Thus, we are unable to determine the relationship between p factor scores and the intrinsic functional connectivity of the cerebellum. We anticipate that current state-of-the-art multiband image acquisition protocols will routinely allow for full coverage of the cerebellum and, subsequently, direct analyses of how its intrinsic connectivity may scale as a function of p factor scores. The observational nature of our study represents another limitation as we cannot establish causal links between p factor scores and intrinsic connectivity. Longitudinal designs may better address causality and temporal order of these phenomena. Future research employing transcranial magnetic stimulation, closed-loop fMRI, and intervention designs can further map causal relationships.

These limitations notwithstanding, our current work provides initial evidence for unique connectome wide functional signatures of the p factor. Consistent with emerging transdiagnostic and dimensional research into the neural basis of psychopathology(11, 12, 44), our analyses reveal that increased broad risk for all common forms of mental illness is associated with higher intrinsic connectivity between visual association cortex and both frontoparietal and default mode networks. Such hyper-connectivity suggests that increased risk for psychopathology may be manifest as greater effortful or less efficient executive control as well as poor regulation of self-referential information processing. These patterns place alterations of the functional connectome squarely in the middle of converging theories of network dysfunction in psychopathology, further suggesting the p factor as a promising tool in clinical neuroscience.

## Acknowledgements and Disclosures

We thank the Duke Neurogenetics Study participants and the staff of the Laboratory of NeuroGenetics. The Duke Neurogenetics Study received support from Duke University as well as US-National Institutes of Health grants R01DA033369 and R01DA031579. ARK, and ARH received further support from US-National Institutes of Health grant R01AG049789. MLE was supported by the National Science Foundation Graduate Research Fellowship under Grant No. NSF DGE-1644868. The authors declare no competing financial interests.

