## Supplementary Materials for "A Connectome Wide Functional Signature of Transdiagnostic Risk for Mental Illness"

### Supplemental Information

#### Supplemental Methods

##### Measuring Psychiatric Symptoms

We assessed symptoms from 11 different psychiatric disorders using both the electronic Mini International Neuropsychiatric Interview (e-M.I.N.I.) and self-report questionnaires. The e-M.I.N.I. is a short, structured diagnostic interview for DSM-IV and ICD-10 psychiatric disorders(1). Clinical psychologists, graduate students, and post-baccalaureate research assistants under the supervision of a licensed clinical psychologist conducted interviews.

P-factors were originally derived in 1250 subjects of the Duke Neurogenetic Sample (DNS) (including all subjects who completed resting-state fMRI), 250 (20%) of these participants met criteria for at least one Axis I or II disorder, including 136 with alcohol use disorders, 48 with non-alcohol substance use disorders, 59 with major depressive disorders, 35 with bipolar disorders, 22 with panic disorder (no agoraphobia), 21 with panic disorder including agoraphobia, 11 with social anxiety disorder, 22 with generalized anxiety disorder, 15 with obsessive-compulsive disorder, 11 with eating disorders, 2 with post-traumatic stress disorder, and 132 with at least one comorbid diagnosis.

*Internalizing Symptoms.* Anxiety and depressive symptoms were assessed with the 62-item Mood and Anxiety Symptom Questionnaire—Short Form (MASQ-SF), the 20-item State-Trait Anxiety Inventory—Trait (STAI-T), the 20-item Center for Epidemiological Studies on Depression scale (CESD), and symptom counts of panic disorder, agoraphobia, and social phobia from the e-M.I.N.I. The MASQ-SF is a well-validated measure(2) yielding four subscales assessing symptoms experienced within the last seven days specific to Anxious Arousal, General

Distress Anxiety, Anhedonic Depression, and General Distress Depression. The STAI-T was used to assess participants' general tendency to perceive situations as threatening and to respond to such situations with subjective feelings of apprehension and tension(3). The CESD was used to assess depressive symptoms within the past week(4).

Using these measures, five scores of anxiety and depressive symptoms were created: 1) a MASQ-SF anxiety score was created by standardizing (z-scoring) and then averaging the Anxious Arousal and General Distress scales; 2) the sum total score on the STAI-T self-report questionnaire was used as a second measure of trait anxiety; 3) a MASQ-SF depression score was created by z-scoring and averaging the Anhedonic Depression and General Distress Depression scales; 4) the sum total score on the CESD scale was used as a second measure of depression; 5) e-M.I.N.I. symptom counts of social phobia, panic disorder, and agoraphobia were z-scored and then averaged to create a count of fears/phobias symptoms.

*Externalizing Symptoms.* Antisocial personality/psychopathy, delinquency, and substance abuse and dependence symptoms were assessed using the 29-item Self Report of Psychopathy—Short Form scale, the 49-item Self Report of Delinquency scale (revised), the 10-item Alcohol Use Disorders Identification Test, the 13-item Recreational Drug Use questionnaire, and symptom counts of cannabis abuse and dependence from the e-M.I.N.I. We did not include nicotine dependence symptoms in our assessment of externalizing disorders given that less than 2% of our sample reported ever smoking cigarettes. The Self Report of Psychopathy scale assesses the Interpersonal, Affective, Lifestyle, and Antisocial factors of psychopathy(5). The Self Report of Delinquency scale assesses the frequency with which individuals have engaged in aggressive and delinquent behavior, alcohol and drug use, and related offenses(6). The Alcohol Use Disorders Identification Test assesses the frequency with which participants report hazardous and harmful use of alcohol as well as alcohol dependence(7). The Recreational Drug Use scale assesses the frequency with which participants report using other substances (e.g., cocaine) in their lifetime.

Using these measures, five scores of antisocial personality/psychopathy and substance abuse and dependence symptoms were created: 1) the Self Report of Psychopathy was used to measure antisocial personality and psychopathy symptoms; 2) the Self Report of Delinquency was used to measure delinquent symptoms; 3) alcohol abuse and dependence symptoms were measured using the Alcohol Use Disorders Identification Test total score; 4) cannabis abuse and dependence symptoms were measured using a symptom count from the e-M.I.N.I.; and 5) other substance use and abuse were assessed using the Recreational Drug Use total score.

*Thought Disorder Symptoms.* Three scores of obsessive-compulsive disorder, mania, and psychosis were created using symptom counts from the e-M.I.N.I. Mania included counts of both manic and hypomanic symptoms (see Supplemental Table 1 for inter-correlations among psychiatric symptoms).

*Structure of Psychopathology.* We used confirmatory factor analysis to fit three standard models: correlated factors, bi-factor, and one-factor models(8, 9). All confirmatory factor analyses were performed in MPlus version 7.4(10) using the weighted least squares means and variance adjusted (WLSMV) algorithm. The WLSMV estimator is appropriate for categorical and nonmultivariate normal data and provides consistent estimates when data are missing at random with respect to covariates(11). We assessed how well each model fit the data using the chi-square value, the comparative fit index (CFI), the Tucker-Lewis index (TLI), and the root-mean square error of approximation (RMSEA). Nonsignificant chi-square tests indicate good model fit; nonetheless, this test is generally overpowered in large sample sizes such as ours. CFI and TLI values greater than 0.90 indicate adequate fit; RMSEA scores less than .08 are considered acceptable(12). Supplemental Table 2 shows all three models (correlated factors, bi-factor, and one-factor) with standardized factor loadings and the correlations between the three specific factors.

*Correlated Factors Model.* Using this model, we tested the hypothesis that there are latent trait factors, each of which influences a subset of the diagnostic symptoms. In our case, we tested three factors representing Externalizing (with loadings from alcohol, cannabis, other drugs, and antisocial personality disorder/psychopathy, and delinquency), Internalizing (with loadings from MASQ-SF depression, CESD, MASQ-SF anxiety, STAI-T, and fears/phobias), and Thought Disorder (with loadings from obsessive-compulsive disorder, mania, and psychosis). The model allows the Externalizing, Internalizing, and Thought Disorder factors to be correlated.

We found that the model provided a moderately adequate fit to the data: $\chi^2(62, N = 1,246) = 620.813$, CFI = 0.903, TLI = 0.878, RMSEA =.085, 90% confidence interval (CI) = [.079, .091]. Loadings on the three factors were all positive and statistically significant (all $p < .001$). Correlations between the three factors were all positive and ranged from .257 between Internalizing and Externalizing to .415 between Internalizing and Thought Disorder (see Supplemental Table 2).

*Bi-Factor Model.* Using this model, we tested the hypothesis that the symptom measures reflect both the general ‘p factor’ and three specific forms of psychopathology that are orthogonal to p. For example, depression symptoms loaded on both the ‘p factor’ and on the Internalizing factor. The specific factors represented the constructs of Externalizing, Internalizing, and Thought Disorder.

This model had a Heywood case, an estimated variance that was negative for one of the lower-order symptom factors (specifically, fears/phobias), suggesting this was not a valid model.

Inspection of the results revealed the source of the convergence problem. We respecified the model to be consistent with a previous model in which Thought Disorders were subsumed in p(13). In this model, p served as a general factor and Internalizing and Externalizing factors serving as additional unique sources of variation apart from p (see Supplemental Table 2). This revised model fit the data well: $\chi^2$ (55, N = 1,246) = 385.084, CFI = 0.943, TLI = 0.919, RMSEA = .069, 90% CI [.063, .076]. Loadings on the ‘p factor’ were all positive and statistically significant (all $p < .05$). The highest standardized loadings were for mania (0.625) and MASQ anxiety (0.517). Similarly, the loadings for the two specific factors were all positive and statistically significant (all $p < .001$).

*One Factor Model.* Using this model, we tested whether the specific factors are needed in a simple structural model that assigned each diagnostic symptom only to the ‘p factor.’ Loadings on p were all positive and statistically significant (all $p < .001$) (see Supplemental Table 2). However, this model did not fit the data well: $\chi^2$(65, N = 1,246) = 2696.079, CFI = 0.544, TLI = 0.452, RMSEA = .180, 90% CI [.174, .186].

#### Supplemental Results

##### Connectome Wide Association study (CWAS) with Power 2011 atlas

To ensure the main CWAS results were independent of parcellation scheme, we redid the CWAS analysis with another commonly used resting state atlas(14). Like our resting state data, this atlas was defined in the MNI space in the volume and included subcortical and cortical regions of interest ROIs. We removed 8 of the 264 ROIs because they were in the cerebellum and our resting-state data lacked adequate coverage of the cerebellum. Average resting-state time courses were extracted from 5mm spheres around each of the remaining 256 coordinates in the atlas and entered into the CWAS.

In a parallel analysis to the Lausanne atlas described in the main text, seed-based connectivity analysis was conducted to generate a whole-brain functional connectivity map for each participant. Then, the average distance (1 minus the Pearson correlation) between each pair of participant's functional connectivity maps is computed, resulting in a distance matrix encoding the multivariate similarity between each participant's connectivity map. Finally, multi-dimensional matrix regression (MDMR) is used to generate a pseudo-F statistic quantifying the strength of the association between the phenotype of interest, here p factor scores, and the distance matrix created in the second step, controlling for the sex covariate.

Of the 256 ROIs investigated, multi-dimensional matrix regression (MDMR) analysis revealed one regions with whole-brain connectivity patterns significantly associated with p factor scores in the left lingual gyrus. Of note, this ROI is very close all three of significant left hemisphere ROIs found in the original CWAS with the Lausanne ROIs (see figure S2).

**Controlling for parental socioeconomic background**

We asked participants to report their biological, step-, or guardian mother's and father's highest education level before they turned 18 years old, on a scale from 1, "No high school," to 10, "MD/PhD/JD/PharmD.". This variable was than recoded to a 1-4 scale (1 = no high school diploma, 2 = completed high school, 3 = completed a bachelor's degree, 4 = received a graduate degree). We took the maximum of their mother's or father's education level to create one measure of parental education level (mean: 3.51, sd = .73; .7% never received a high school diploma, 12.1 % completed high school, 23.2% completed bachelor's degree and 63.9% received a graduate degree). Maximum parental education was included as a covariate in a CWAS analyses of the 'p factor' scores along with sex. The substantive results of this analysis were identical. The exact same visual seeds survived multiple correction in the first level of the CWAS.

**Controlling for Diagnoses.**

As described above, psychiatric or substance use diagnosis was assessed using the e-M.I.N.I. and self-report questionnaires. 133 individuals in our final resting-state sample had at least one diagnoses. The presence of diagnosis was turned into a binary covariate and included in a CWAS analysis of the p-factor along with sex. The substantive results of this analysis were identical. The exact same visual seeds survived multiple correction in the first level of the CWAS.

Table 1. *Inter-correlations among Individual Psychiatric Symptoms.*

| | Psychosis (1) | Mania (2) | OCD (3) | Fears (4) | MASQ Depression (5) | CESD (6) | MASQ Anxiety (7) | STAI (8) | Psychopathy (9) | Delinquency (10) | Cannabis (11) | Alcohol (12) | Drug Use (13) |
|---|---|---|---|---|---|---|---|---|---|---|---|---|---|
| 1 | 1 | | | | | | | | | | | | |
| 2 | .072 | 1 | | | | | | | | | | | |
| 3 | **.262** | **.490** | 1 | | | | | | | | | | |
| 4 | **.154** | **.293** | **.431** | 1 | | | | | | | | | |
| 5 | **.147** | **.219** | **.199** | **.266** | 1 | | | | | | | | |
| 6 | .114 | **.256** | **.225** | **.244** | **.842** | 1 | | | | | | | |
| 7 | **.129** | **.172** | **.183** | **.240** | **.565** | **.609** | 1 | | | | | | |
| 8 | .075 | **.258** | **.250** | **.330** | **.770** | **.727** | **.530** | 1 | | | | | |
| 9 | **.193** | **.286** | .058 | .014 | **.193** | **.235** | **.267** | **.209** | 1 | | | | |
| 10 | .019 | **.239** | **.117** | .033 | **.135** | **.194** | **.245** | **.130** | **.531** | 1 | | | |
| 11 | **.158** | **.408** | **.242** | **.201** | **.137** | **.142** | **.143** | **.202** | **.368** | **.381** | 1 | | |
| 12 | .073 | **.149** | -.036 | -.006 | .033 | .055 | **.155** | .037 | **.373** | **.585** | **.452** | 1 | |
| 13 | .038 | **.240** | .061 | -.008 | .035 | .030 | .062 | .047 | **.357** | **.590** | **.714** | **.566** | 1 |

*Note.* Correlations with $p < .01$ are shown in bold. OCD = obsessive-compulsive disorder; MASQ = Mood and Anxiety Symptom Questionnaire; CESD = Center for Epidemiological Studies – Depression Scale; STAI = State-Trait Anxiety Inventory – Trait Anxiety Scale.

Table 2. *Model Fit Statistics, Standardized Factor Loadings, and Factor Correlations From Three Different Confirmatory Factor Models.*

| | Correlated Factors Model | | | | Bi-factor Model | | | | One Factor Model | |
|---|---|---|---|---|---|---|---|---|---|---|
| Statistics, Loadings, and Correlations | Model Fit | THT | INT | EXT | Model Fit | p | INT | EXT | Model Fit | p |
| Statistic | | | | | | | | | | |
| Chi-Square (WLSMV) | 620.813 | | | | 385.084 | | | | 2696.079 | |
| Degrees of Freedom | 62 | | | | 55 | | | | 65 | |
| Comparative Fit Index | .903 | | | | .943 | | | | .544 | |
| Tucker-Lewis Index | .878 | | | | .919 | | | | .452 | |
| RMSEA [90% CI] | .085 [.079, .091 | | | | .069 [.063, .076] | | | | .180 [.174, .186] | |
| Standardized Factor Loadings | | | | | | | | | | |
| Psychosis | | .338 | | | | .307 | | | | .197 |
| Mania | | .786 | | | | .625 | | | | .434 |
| OCD | | .595 | | | | .499 | | | | .325 |
| Fears | | | .343 | | | .303 | .193 | | | .299 |
| MASQ Depression | | | .859 | | | .358 | .858 | | | .553 |
| CESD | | | .892 | | | .443 | .790 | | | .548 |
| MASQ Anxiety | | | .701 | | | .517 | .457 | | | .512 |
| STAI | | | .811 | | | .406 | .715 | | | .521 |
| Psychopathy | | | | .640 | | .491 | | .410 | | .535 |
| Delinquency | | | | .782 | | .404 | | .644 | | .719 |
| Cannabis | | | | .569 | | .300 | | .498 | | .566 |
| Alcohol | | | | .703 | | .206 | | .718 | | .563 |
| Drug Use | | | | .749 | | .150 | | .822 | | .664 |
| Factor Correlation | | | | | | | | | | |
| INT | | .652 | | | | | | | | |
| EXT | | .556 | .249 | | | | | | | |

*Note.* THT = Thought Disorders factor, INT = Internalizing factor; EXT = Externalizing Disorders factor; p = ‘p factor’; OCD = obsessive-compulsive disorder; MASQ = Mood and Anxiety Symptom Questionnaire; CESD = Center for Epidemiological Studies – Depression Scale; STAI = State-Trait Anxiety Inventory – Trait Anxiety Scale.

**Supplementary Figure S1.** Distribution of p factor in our final resting-state sample. While individuals with a diagnosed mental illness have higher P-factor scores on average, their distribution is continuous with the overall sample. Individuals with a diagnosis span the whole range of p-factor values and largely overlap with the distribution of individuals without a diagnosis.

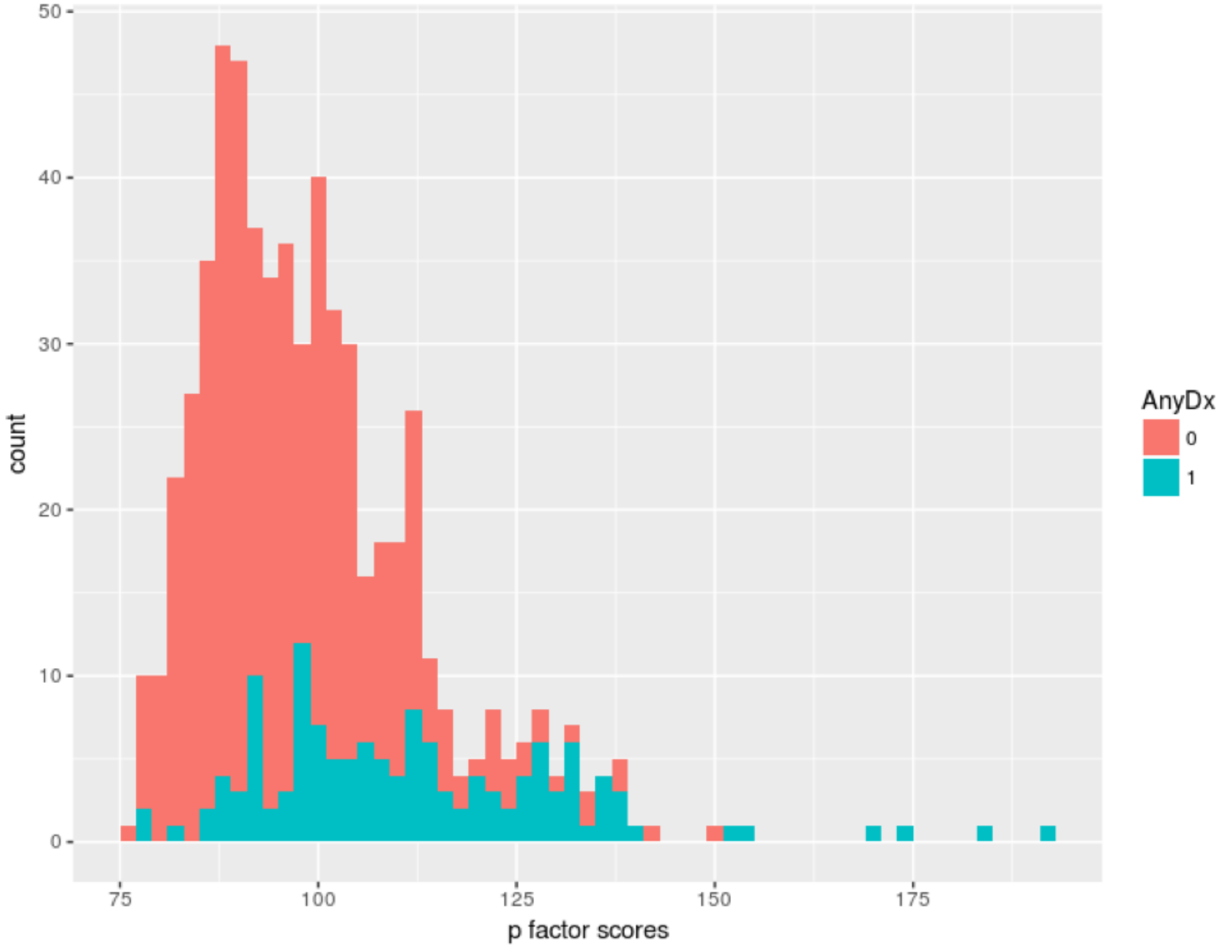

**Supplementary Figure S2.** Follow-up connectivity analyses of the seed identified in the Power 264 parcellation. MDMR reveals a highly consistent pattern of connectivity that closely follows the results in the main text with the Lausanne atlas. All results were projected from the volume onto a surface to aid visualization. Left panel: MDMR-derived seed regions. Middle panel: average intrinsic connectivity for each seed. Right panel: connectome wide intrinsic connectivity patterns for each seed as a function of p factor scores.

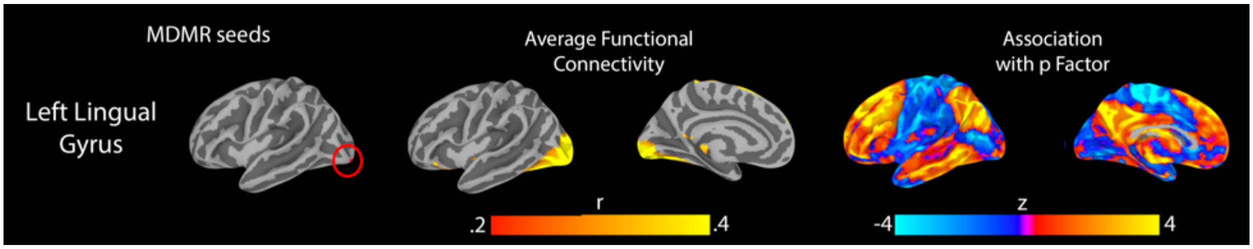
